# Increased Occurrence of *Treponema spp*. and Double-species Infections in Patients with Alzheimer’s Disease

**DOI:** 10.1101/2021.11.04.467230

**Authors:** Michal Nemergut, Tereza Batkova, Dana Vigasova, Milan Bartos, Martina Hlozankova, Andrea Schenkmayerova, Barbora Liskova, Katerina Sheardova, Martin Vyhnalek, Jakub Hort, Jan Laczo, Ingrid Kovacova, Michal Sitina, Radoslav Matej, Radim Jancalek, Martin Marek, Jiri Damborsky

## Abstract

**Objective:** Although the link between microbial infections and Alzheimer’s disease (AD) has been demonstrated in multiple studies, the involvement of pathogens in the development of AD remains unclear. Therefore, this theory beckons further systematic investigation. In this study, we have examined the association between the 10 most widely discussed viral and bacterial pathogens found in serum and cerebrospinal fluid (CSF) from patients with AD.

**Methods:** We have used an *in-house* developed multiplex PCR kit for simultaneous detection of five bacterial and five viral pathogens in serum and CSF from 50 AD patients and 53 healthy controls. Data analysis was performed with multiple statistical methods: Fisher’s exact test, chisquare goodness of fit test, and one-sample proportion test.

**Results:** We observed an increased frequency of AD patients tested positive for *Treponema spp.* (AD: 62.2%; CTRL: 30.3%; *p*-value = 0.007). Furthermore, we confirmed a significantly higher prevalence of cases with two and more simultaneous infections in AD patients compared to controls (AD: 24%; CTRL 7.5%; *p*-value = 0.029). The studied pathogens were widespread equally in serum and CSF. *Borrelia burgdorferi*, human herpesvirus 7, and human cytomegalovirus were not detected in any of the studied samples.

**Discussion:** An increased prevalence of *Treponema* spp. and double-species infections in AD patients compared to the healthy controls provides further evidence of the association between microbial infections and AD. Paralleled analysis of multiple sample specimens provides complementary information and is advisable for future studies.

## Introduction

Alzheimer’s disease (AD) is an irreversible, progressive neurodegenerative pathology. It accounts for 60-80% of dementia [1], a general term for memory loss and other cognitive abilities, serious enough to interfere with daily life. The AD aetiology is largely unknown to this day. The most widely accepted hypothesis for AD pathogenesis is the amyloid cascade hypothesis, which states that the cause of AD is a senile plaque formation by the β-amyloid peptide and the generation of neurofibrillary tangles of hyperphosphorylated tau protein [2]. However, the reason for the initial accumulation of β-amyloid is unknown in most patients with sporadic AD [3]. Previous research brought some evidence about the role of infection in AD pathophysiology [4]. AD was linked to infectious agents in the brain tissue, specifically to herpes simplex virus 1 (HSV-1), almost 30 years ago [5]. Ever since then, a growing body of evidence has associated the infection of various other viruses, including human herpesvirus 6 and 7 (HHV-6 and HHV-7) [6–9], human cytomegalovirus (CMV) [9–12] and Epstein-Barr virus (EBV) [7] with the risk of developing AD. Different bacterial species, including *Chlamydia pneumoniae* [13], *Helicobacter pylori* [14,15], *Borrelia burgdorferi* [16], *Porphyromonas gingivalis* [17] and *Treponema spp.* [18] were also implicated in AD pathogenesis.

However, the data driving the infectious hypothesis of AD has been conflicting, causing an inevitable controversy in the field [19]. Although the extent of causal contribution of infections is not conclusive, certain microorganisms could act as accelerants, exacerbating the disease once it is established [19]. Evidence suggests that severe sepsis survivors are more likely to develop substantial and persistent cognitive impairment and functional disability [20]. Moreover, it is unclear whether the disease process involves a single microorganism or several species, acting independently or in combination [19]. Therefore, more research in this field is needed, comparing the data collected from multiple specimen types and involving alternative methods for pathogen detection.

In this study, we developed a new multiplex PCR kit for simultaneous detection of acute inflammation of the 10 most prevalent bacterial and viral pathogens associated with AD. Based on this, we compared the occurrence of single and multiple infections in AD patients and controls. In addition, we compared the prevalence of the studied pathogens between serum and CSF matched AD patients and controls.

## Material and methods

### Study participants

We have analysed samples from 50 living patients diagnosed with AD by biomarkers in mild cognitive impairment (MCI) or dementia stages (mean age 71 years; 17 males and 33 females) recruited from the Czech Brain Aging Study (CBAS) [21]. They consisted of 5 serum samples and 45 serum and CSF matched samples. All CBAS patients underwent the complex diagnostic process including neuropsychological examination, brain MRI, neurological examination, and routine blood tests. Control samples were obtained from 53 donors (mean age 45 years; 19 males and 34 females). Serum and CSF samples were obtained from 18 CBAS subjects and 27 cognitively healthy subjects undergoing spinal tap for differential diagnosis of headache or facial palsy. They consisted of 9 serum samples, 3 CSF samples and 33 matched CSF and serum samples. To exclude AD pathology, subjects were selected with the following criteria: the subjects had negative AD biomarkers in CSF or were cognitively stable for 3 years of follow up and APOE 4 negative. In the cognitively healthy headache and facial palsy group, only subjects with physiological parameters of CSF were included. Eight brain samples were obtained from the patients undergoing surgery for pharmacoresistant epilepsy. The name of Ethics Committee is FN u sv. Anny v Brne and the approval numbers are 45V/2016 and 46V/2016-AM. All procedures were performed in consistence with Good clinical practice and after signing informed consent.

### DNA isolation and PCR primers design

Samples were thawed and cut into smaller pieces in case of brain tissue. DNA was extracted using RTP Pathogen Kit (STRATEC, USA). Each extract was tested for all ten pathogens by amplifying species-specific gene sequences using species-selective primers. All primers used in this study were designed in-house using the Primer-BLAST tool for finding primers specific to the polymerase chain reaction (PCR) template (Table 1).

**Table 1.**
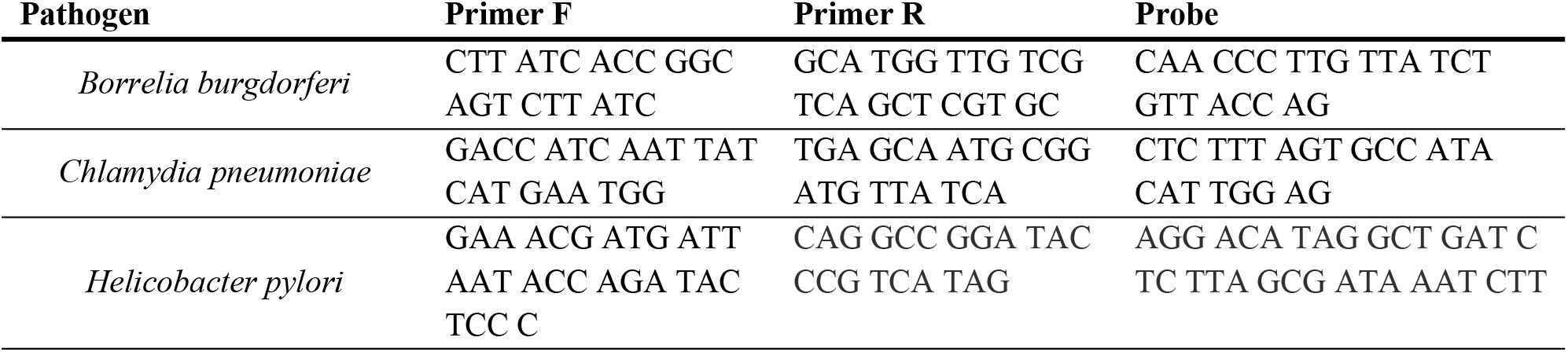
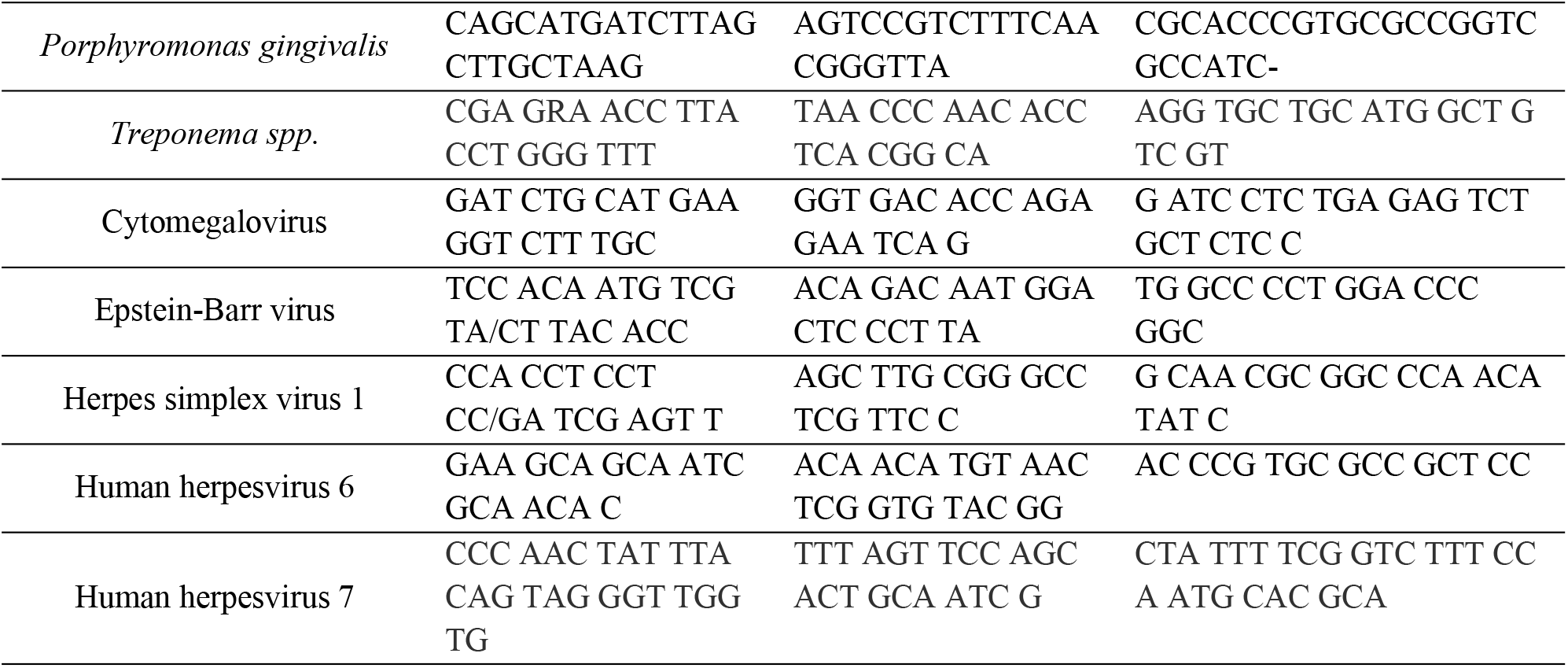
List of primers (5’-3’) used in the polymerase chain reaction.

### Polymerase chain reaction

Multiplex PCR for detecting five bacterial species *Chlamydia pneumoniae*, *Helicobacter pylori*, *Borrelia burgdorferi*, *Porphyromonas gingivalis*, *Treponema spp.,* and five viral species HSV-1, EBV, HHV-6, HHV-7, CMV, was developed. The PCR MasterMixes of 20 μl contained 2U of Taq polymerase (Thermofisher Scientific, United Kingdom) and 2U of Uracil-D-glycosylase (New England Biolabs, USA). Conditions of PCR were as follows: incubation for 2 min at 37°C to perform deactivation of old amplicons by Uracil-D-glycosylase, activation for 15 min at 95°C, 45 cycles of denaturation, each 5 s at 95°C, annealing for 40 s at 60°C and polymerisation for 20 s at 72°C. A positive signal was read in the FAM channel and a signal for internal control in the HEX/JOE channel. Each PCR experiment included either AD or control DNA, a positive DNA template, and a no-DNA control. A subject was considered positive for a given species if the extract contained a positive signal with Ct ≤ 35.0. Positive controls were performed using plasmid vectors carrying loci for respective pathogens. Reagents and disposable supplies were examined for DNA as the negative controls. All tests were successfully validated using international panels available on a regular basis from INSTAND, e.V. (Germany).

### Statistical analysis

Statistical methods used for data analysis included Fisher’s exact test, chi-square goodness of fit test and one-sample proportion test. The significance of the difference between AD patients and controls was determined by Fisher’s exact test. The chi-square goodness of fit test was used to assess the differences in the distribution of the prevalence of cases without infection, with single infection and multiple infections. Differences between individual groups were further analysed using a one-sample proportion test with Bonferroni correction for multiple testing. All statistical analyses were performed using the software R (version 4.0.4). *P*-values less than 0.05 were considered statistically significant.

## Results

### Comparison of bacterial and viral infections between AD patients and controls

To investigate the possible association between increased bacterial/viral infections and AD, we divided all AD patients and controls into four groups: *group 1 –* tested positive for at least one pathogen; *group 2* – tested negative for all studied pathogens; *group 3* – tested positive for a single pathogen; *group 4* – simultaneously tested positive for two or more pathogens. The data from 50 AD patients and 53 controls, which were tested positive for at least one bacterial or viral pathogen, are compared in Figure 1.

**Figure 1.**
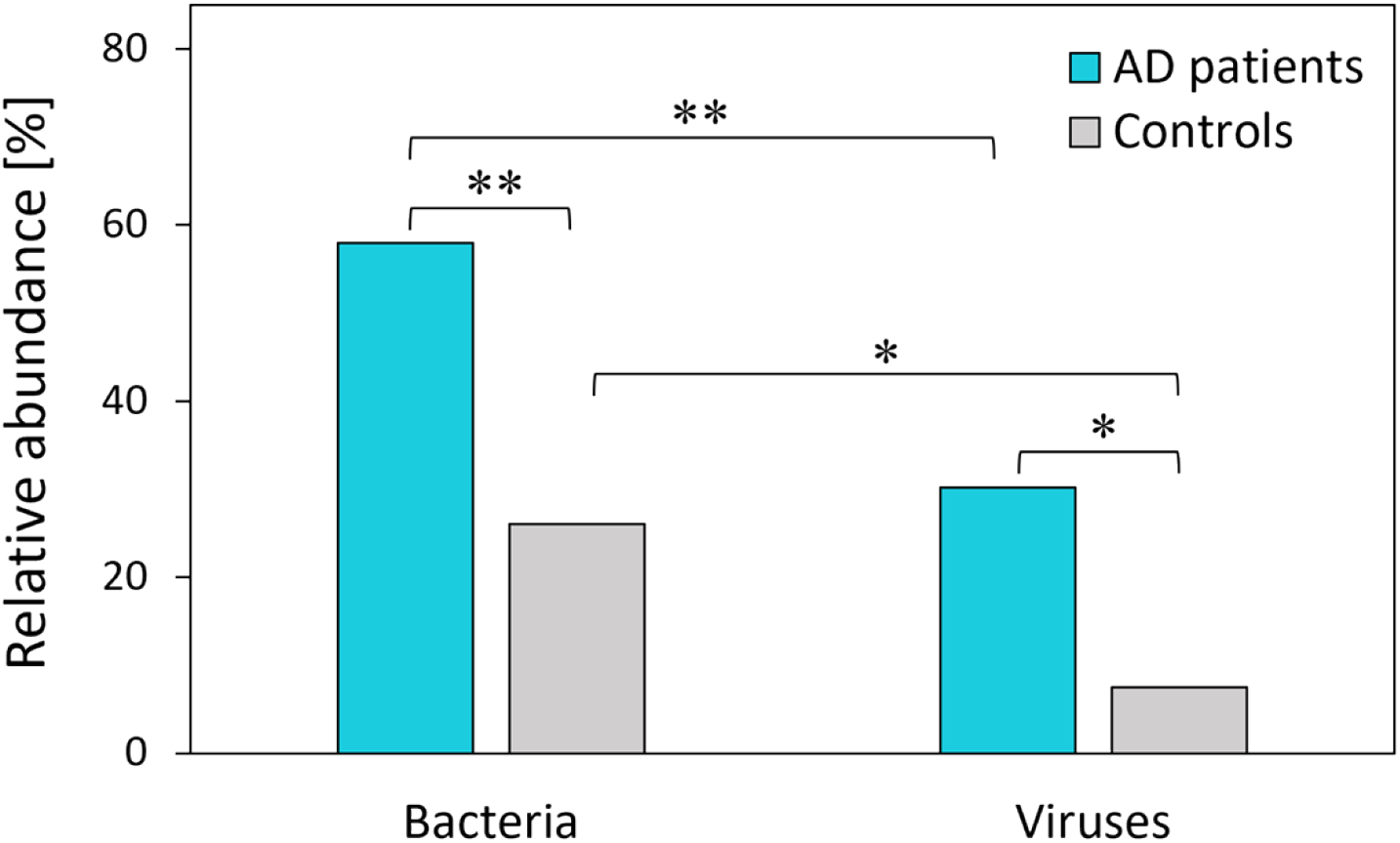
Percentage of AD patients (turquoise) and controls (grey) with at least one bacterial or viral pathogen detected. Asterisks represent statistical significance (* *p*-value ≤ 0.05; ** *p*-value ≤ 0.01).

Our results showed a significant difference between AD patients and controls in the prevalence of both bacterial (*p*-value = 0.006) and viral (*p*-value = 0.016) pathogens (Table 2). In *group 1*, almost twice as many AD patients (58%) were positive for bacterial infection compared to controls (30.2%). Similarly, the frequency of viral pathogens was 3.5 times higher in AD patients (26%) than in controls (7.5%). No significant difference in the prevalence of bacterial and viral pathogens was found in studied groups in terms of gender. A comparison of bacterial and viral prevalence revealed that the bacterial pathogens are present in a significantly higher proportion than viral pathogens in AD patients (*p*-value = 0.006) as well as the control group (*p*-value = 0.016). The prevalence of cases without infection, with single infection and with multiple infections (two and more) is compared in Figure 2.

**Table 2.** Percentages of AD patients and controls tested positive for bacterial or viral infections.

| Pathogens |  | AD patients<br>(n = 50) | Controls<br>(n = 53) | *p*-values |
| --- | --- | --- | --- | --- |
| At least one<br>pathogen | Total | 66 % | 32.1 % | < 0.001 |
|  | Bacteria | 58 % | 30.2 % | 0.006 |
|  | Viruses | 26 % | 7.5 % | 0.016 |
| No<br>pathogen | Total | 34 % | 67.9 % | < 0.001 |
| Single<br>pathogen | Total | 42 % | 24.5 % | 0.093 |
| Multiple<br>pathogens | Total | 24 % | 7.5 % | 0.029 |

**Figure 2.**
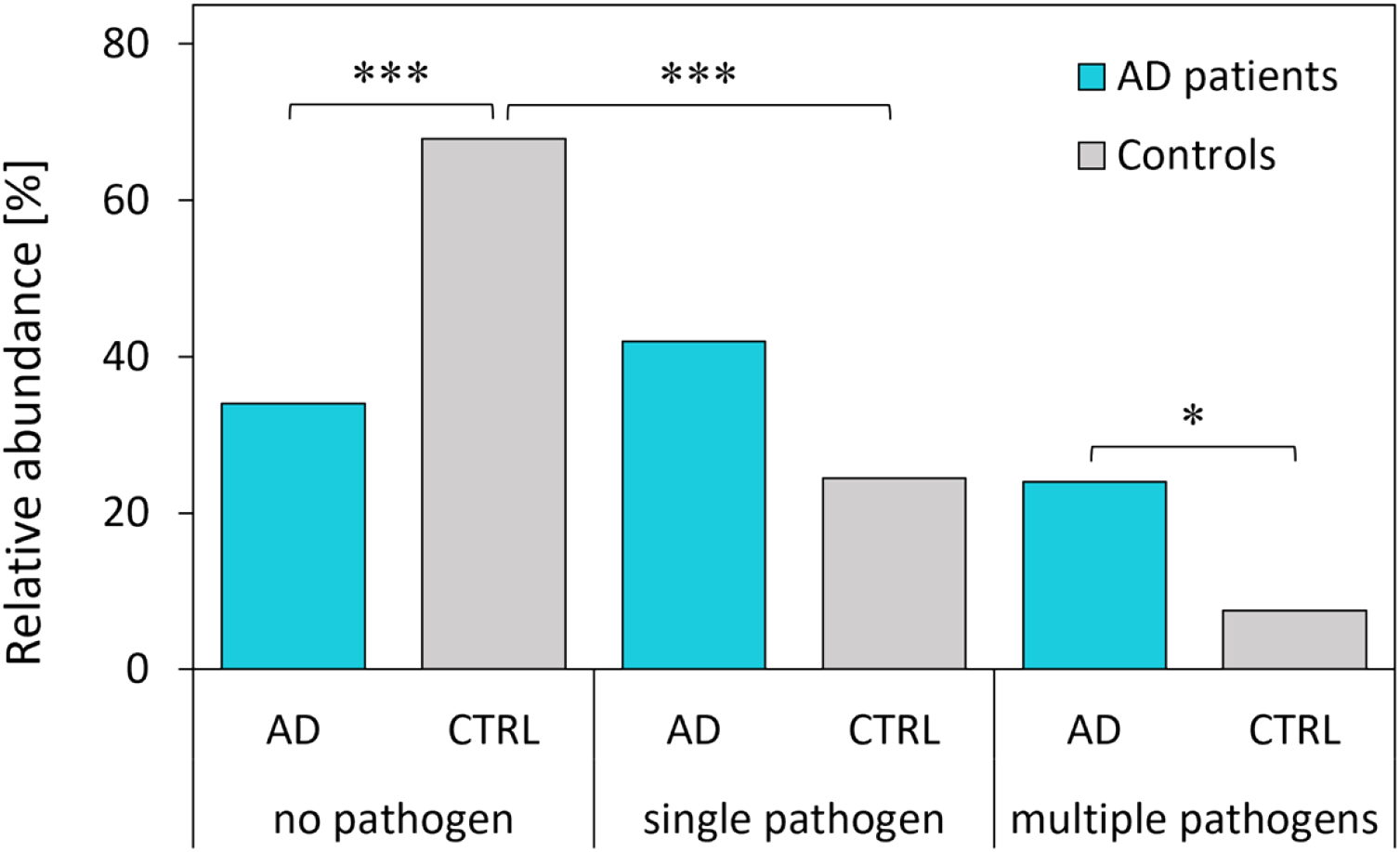
Number of pathogens simultaneously detected in AD patients (turquoise) and controls (grey). Asterisks represent statistical significance (* *p*-value ≤ 0.05; *** p-value ≤ 0.001).

Based on the Chi-square goodness of fit test, the distribution of the groups is equal among AD patients (*p*-value = 0.295) while it is unequal among the controls (*p*-value < 0.001). The most prevalent group among the controls is the one that tested negative for all studied pathogens (Table 2). As expected, the controls without any pathogen represent 67.9% of all controls, and their prevalence is significantly higher compared to 34% of AD patients in the same group (*p*-value ≤ 0.001). The number of AD patients in *group 3* (42%) exceeds the number of controls (24.5%) infected with a single bacterial or viral pathogen; however, this difference is not statistically significant (*p*-value = 0.093). Interestingly, simultaneous infection by two and more different pathogens was detected in 24% of AD patients. This represents a significant prevalence compared to controls where the multiple infections were detected only in 7.5% of controls (*p*-value = 0.029).

### Comparison of the prevalence of studied pathogens in serum and CSF

To explore the prevalence of studied viral and bacterial pathogens in different specimen types, we compared serum, and CSF matched AD patients and controls. The data from 45 AD patients and 33 controls, whoch were tested positive for at least one bacterial or viral pathogen, are compared in Figure 3.

**Figure 3.**
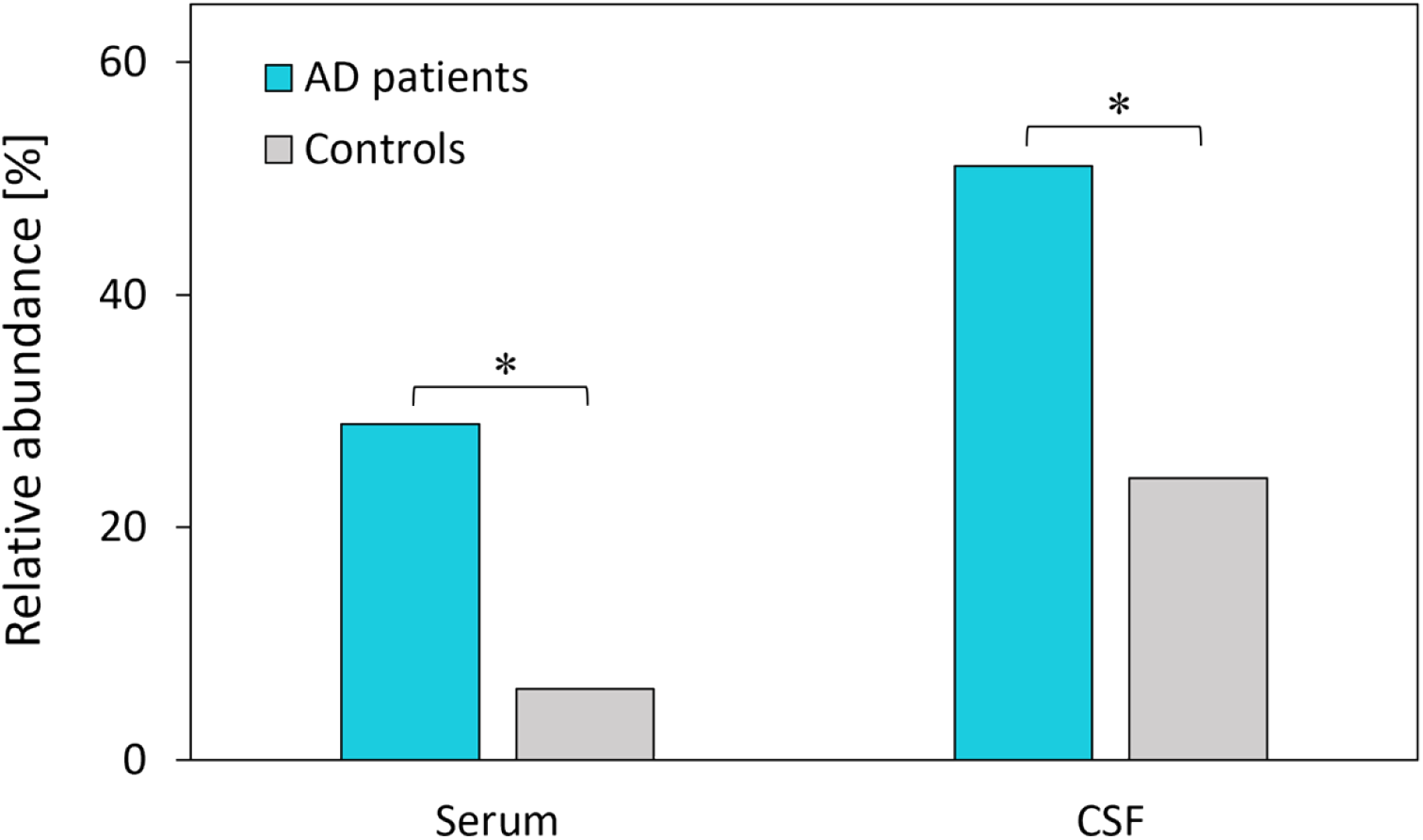
Comparison of serum and CSF matched AD patients (turquoise) and controls (grey) tested positive for at least one pathogen categorized by specimen type. Asterisks represent statistical significance (* *p*-value ≤ 0.05).

Similarly, as in the case with all studied subjects, the results showed a significantly higher prevalence of pathogens among AD patients than controls (Table 3). The pathogens were found with a higher frequency in CSF than in serum; however, this difference is not considered statistically significant (AD: *p*-value = 0.052; CTRL: *p*-value = 0.082). Among AD patients, 51.1% had at least one pathogen in CSF, 28.9% had at least one pathogen in serum, and 17.7% had at least one pathogen in both CSF and serum. In the control group, 24.2% of controls were tested positive for at least one pathogen in CSF, and 6.1% were tested positive for at least one pathogen in serum. None of the controls was tested positive for any pathogen in CSF and serum at the same time.

**Table 3.** Percentages of AD patients and controls tested positive for at least one pathogen in serum and CSF.

| Specimen type | AD patients (n = 45) | Controls (n = 33) | $p$-value |
| --- | --- | --- | --- |
| Serum | 28.9 % | 6.1 % | 0.018 |
| CSF | 51.1 % | 24.2 % | 0.020 |
| $p$-value | 0.052 | 0.082 | |

The relative prevalence of individual bacteria and viruses detected in serum and CSF matched AD patients and controls is shown in Figure 4.

**Figure 4.**
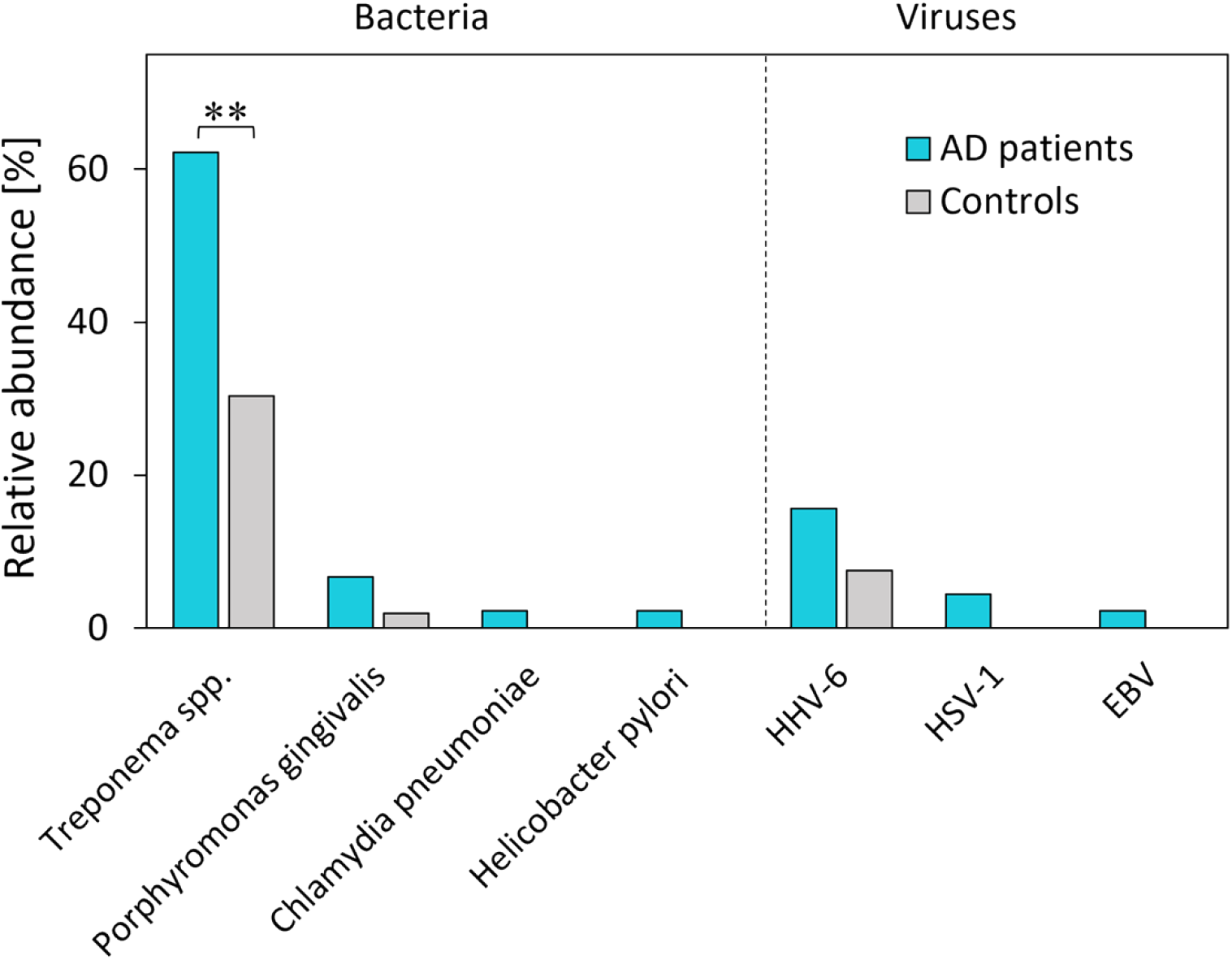
Prevalence of bacterial and viral pathogens between AD patients (turquoise) and controls (grey). Asterisks represent statistical significance (** *p*-value ≤ 0.01).

Our results showed that *Treponema spp.* is the most prevalent pathogen among AD patients (62.2%) as well as controls (30.3%). Moreover, it is the only pathogen found in significantly higher prevalence in AD patients compared to controls (*p* = 0.007). Although significantly less prevalent than *Treponema spp.*, the next two pathogens detected in both AD patients and controls are *Porphyromonas gingivalis* and HHV-6. *Porphyromonas gingivalis* was detected in 6.7% of AD patients and 3.0% of controls (*p*-value = 0.634). HHV-6 represents the most prevalent viral pathogen, however, its prevalence among AD patients (15.6%) compared to controls (6.1%) is not significantly different (*p*-value = 0.062). *Borrelia burgdorferi*, HHV-7, and CMV were detected neither in AD patients nor in the control group. The prevalence of all studied pathogens among AD patients and controls with corresponding *p*-values is listed in Table 4.

**Table 4.** The prevalence of individual pathogens among AD patients and controls.

| | | AD patients<br>(n = 45) | Controls<br>(n = 33) | $p$-values |
| --- | --- | --- | --- | --- |
| Bacteria | *Treponema spp.* | 62.2 % | 30.3 % | 0.007 |
|  | *Porphyromonas gingivalis* | 6.7 % | 3.0 % | 0.634 |
|  | *Chlamydia pneumoniae* | 2.2 % | —<sup>a</sup> | > 0.999 |
|  | *Helicobacter pylori* | 2.2 % | —<sup>a</sup> | > 0.999 |
|  | *Borrelia burgdorferi* | —<sup>a</sup> | —<sup>a</sup> | —<sup>b</sup> |
| Viruses | HHV-6 | 15.6 % | 6.1 % | 0.062 |
|  | HSV-1 | 4.4 % | —<sup>a</sup> | 0.506 |
|  | EBV | 2.2 % | —<sup>a</sup> | > 0.999 |
|  | HHV-7 | —<sup>a</sup> | —<sup>a</sup> | —<sup>b</sup> |
|  | CMV | —<sup>a</sup> | —<sup>a</sup> | —<sup>b</sup> |

<sup>a</sup> pathogens were not detected in these samples; <sup>b</sup> not applicable.

Since the occurrence of most studied pathogens is low, there is not a significant difference in the prevalence of any of the studied pathogens in a specific specimen type. Among AD patients, the presence of *Treponema spp.* was confirmed in 40% of cases in CSF and 22.2% of cases in serum (*p* = 0.110). In the control group, *Treponema spp.* was found in 24.2% and 6.1% of CSF and serum samples, respectively (*p* = 0.082). The prevalence of *Porphyromonas gingivalis* was equal among AD patients and controls in both serum (AD: 2.2%; CTRL: 0%) and CSF (AD: 4.4%; CTRL: 3.0%). HHV-6 was detected in a higher frequency in CSF compared to serum in both AD patients and controls. Among AD patients, it was found in 13.3% of CSF and 2.2% of serum samples (*p*-value = 0.110). The prevalence of HHV-6 in the control group was 6.1% in CSF and no case was found in serum (*p*-value = 0.492).

## Discussion

There is accumulating experimental evidence suggesting the connection between microbial infections and the development of AD [22,23]. In the past, most research in this area has focused on individual pathogens, recently reviewed by Sochocka [24]. However, a growing number of research articles (Table 5) suggest that the aetiology of Alzheimer’s disease could be driven by the coinfection of multiple pathogens [25]. On the other hand, almost all these studies used serum pathogen-specific antibodies to identify infection burden. Therefore, it could not be determined, if the pathogen specific-antibodies result from current, past, or chronic infections. In this work, we used an in-house PCR kit which allows for simultaneous detection of five bacterial pathogens (*Borrelia burgdorferi, Chlamydia pneumoniae, Helicobacter pylori, Porphyromonas gingivalis,* and *Treponema spp.*) and five viral pathogens (CMV, EBV, HHV-6, HHV-7, and HSV-1) in serum and CSF samples. Moreover, previously conducted studies targeted only a single specimen type – serum – while parallel analysis of CSF carried out in this study could increase the relevance of collected data.

**Table 5.**
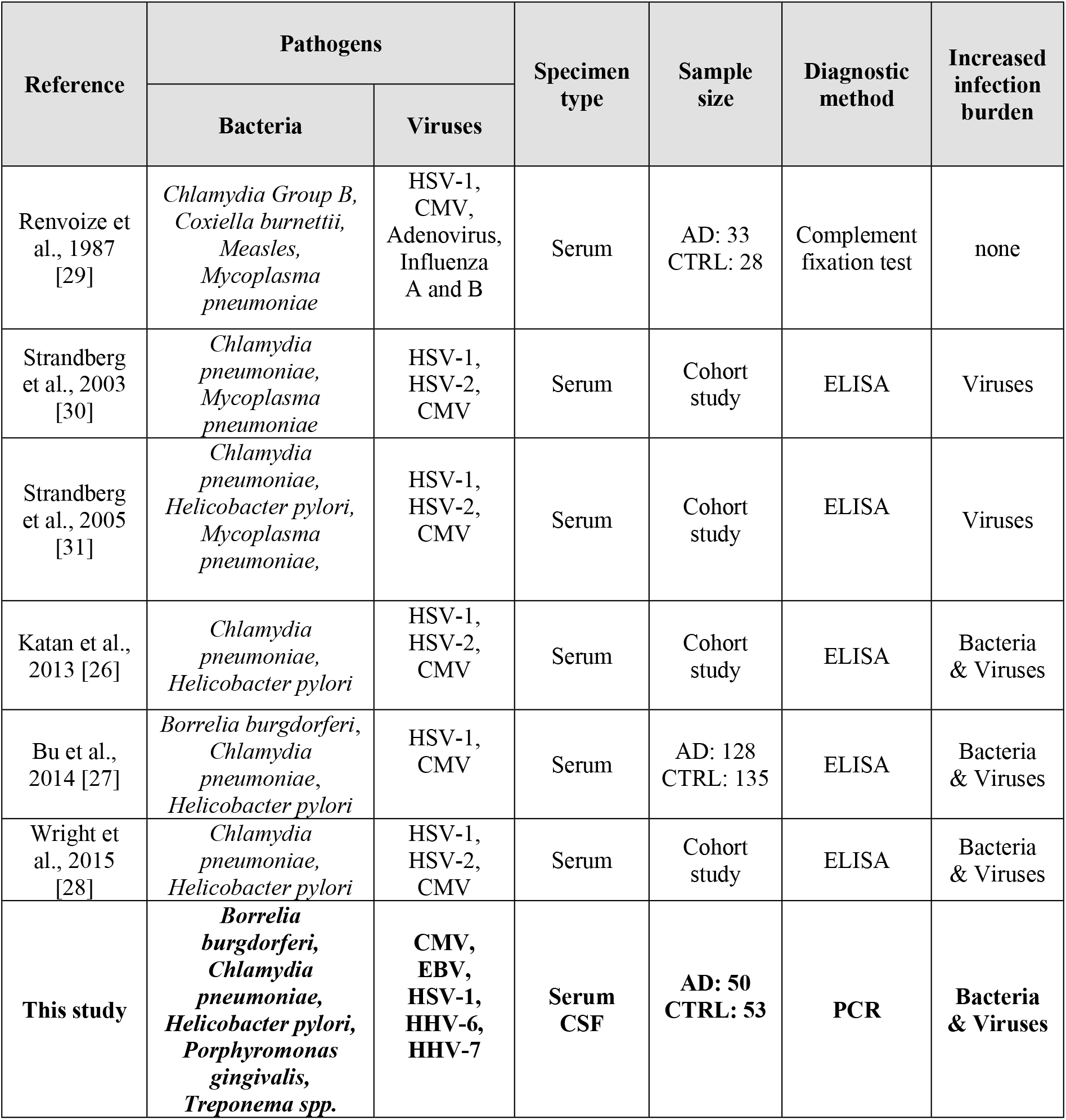
Studies investigating the association of viral and bacterial pathogens with AD and cognitive function (cohort studies).

Our results revealed that both bacteria and viruses are significantly more prevalent in AD-affected samples compared to control samples. This observation is in agreement with recent studies that confirmed an increased bacterial and viral burden in AD patients [26–28]. On the other hand, the cumulative viral and bacterial effect on cognition was not always confirmed [29]. Strandberg et al., who tested seropositivity towards HSV-1, HSV-2, CMV, *Chlamydia pneumoniae*, and *Mycoplasma pneumoniae* in the elderly Finnish population, observed a positive correlation between viral burden and cognitive impairment; however, no association with bacterial burden was observed [30,31]. Conversely, the results of our study showed a significantly higher frequency of bacterial infections than viral infections. This discrepancy can be due to our study’s different and broader range of studied bacteria compared to the studies mentioned above. We observed a significant increase in the prevalence of *Treponema spp.* (*p*-value = 0.082). However, no significant difference in the prevalence was found when comparing the serum and CSF. This observation points to the importance of analysing multiple specimen types in parallel whenever possible.

Interestingly, all tested samples in this study were negative for *Borrelia burgdorferi*, HHV-7 and CMV despite these pathogens having been linked to AD in previous studies [6,10,32–35]. Although, the direct role of CMV in the causation of AD is thought to be unlikely [36]. Our results further confirmed a significantly higher prevalence of cases with multiple (two and more) infections in AD patients compared to controls (AD: 24%; CTRL: 7.5%). Moreover, while the prevalence of AD patients without infection, single infection, and multiple infections is comparable, there is a significant difference in the prevalence of controls without infection and with infection. In addition, a significant difference between AD patients and control tested negative for all studied pathogens (AD: 34%; CTRL: 67.9%) strongly supports a positive correlation between infectious burden and AD. However, the limitation of the present study is the fact that the samples could not be obtained for the age-matched subjects. This is very challenging, particularly for the samples of brains tissues. Further investigations are needed to confirm the impact of microbial infections on AD using the samples for age-matched AD patients and controls.

In summary, to the best of our knowledge, our study is the most comprehensive analysis in terms of detection method, pathogen range, and specimen types conducted thus far. Our results convincingly show that both bacteria and viruses are significantly more prevalent in AD patients than in controls.

## Acknowledgements

The authors would like to express their thanks to the Ministry of Education of the Czech Republic (INBIO - CZ.02.1.01/0.0/0.0/16_026/0008451; ENOCH - CZ.02.1.01/0.0/0.0/16_019/0000868), Technology Agency of the Czech Republic (Permed - TN01000013) and Masaryk University (MUNI/H/1561/2018) for the financial support.

## Author contributions

MN – writing of the manuscript, data analysis, interpretation; TB – testing of samples and data analysis; DV – development of assays, data collection; AS – data analysis; MB – development of assays, data analysis, interpretation, supervision; MH – development of assays, supervision; BL – data collection; KS – a collection of clinical samples, interpretation; MV – a collection of clinical samples, interpretation; JH – a collection of clinical samples, interpretation; JL – a collection of clinical samples, interpretation; IK – statistical analysis; MS – statistical analysis, interpretation; RM – a collection of clinical samples, interpretation; RJ – a collection of clinical samples, interpretation; MM – supervision, funding; JD – experimental design, data interpretation, funding; All co-authors contributed to the revisions of the manuscript.

